# Pre-TCR signaling intensity shapes the TCRβ repertoire

**DOI:** 10.1101/2020.05.04.074922

**Authors:** Elena R. Bovolenta, Eva M. García-Cuesta, Julia Ponomarenko, Mario Mellado, Mario Castro, David Abia, Hisse M. van Santen

## Abstract

Signaling via pre-TCR is essential for initial T cell differentiation. How the signal is triggered and whether its alteration impacts on lymphocyte repertoires is nevertheless debated. We show that a mutation in the transmembrane domain of the CD3ζ chain impairs murine thymocyte development at the double negative (DN) stage coinciding with reduced levels of steady-state pre-TCR signaling. Single particle tracking and TIRF microscopy on mouse primary DN3/DN4 thymocytes shows that wild-type pre-TCR form particles with a broad range of fluorescent intensities whereas mutant complexes are less intensely labeled and move faster, suggesting reduced pre-TCR nanoclustering. Notably, the mutant early-pre-TCR-selected-DP population shows less TCRβ repertoire diversity. We thus propose that the pre-TCR forms stable nanoclusters, the size of which determines signaling efficacy. This signaling does not merely promote maturation of any thymocyte expressing a rearranged TCRβ chain, but shapes the repertoire of DP thymocytes that can audition for positive and negative selection.

## Introduction

T cell differentiation in the thymus occurs through a series of phenotypically well-defined stages. Bone marrow-derived precursors enter the thymus from the blood stream and give rise to double-negative (DN) thymocytes (for lack of expression of the CD4 and CD8 co-receptors). DN thymocytes can be subdivided in four populations based on the expression pattern of the CD44 and CD25 proteins (DN1: CD25^-^CD44^+^; DN2: CD25^+^CD44^+^; DN3: CD25^+^CD44^-^; DN4: CD25^-^CD44^-^), that progressively restrict their lineage potential (Godfrey, Kennedy, Suda, & Zlotnik, 1993; Shah & Zuniga-Pflucker, 2014). DN3 thymocytes rearrange their *Tcrb* locus and only if an in-frame rearrangement is produced, they will express the pre-TCR. The pre-TCR consists of a heterodimer formed by a rearranged TCRβ chain and an invariant pTα chain, non-covalently associated to the signal-transducing CD3γε, CD3δε and CD3ζζ dimers (Groettrup et al., 1993). Successful in-frame rearrangement is checked via a process termed β-selection. This rescues DN3 cells from apoptotic cell death and initiates a proliferative burst concomitant with further differentiation to the CD4^+^CD8^+^ double-positive (DP) stage (von Boehmer, 2005). DP thymocytes rearrange the *Tcra* locus and replace the pTα chain with the TCRα chain. The resulting TCR-expressing thymocytes undergo positive and negative selection, a process that guarantees the generation of a self-MHC restricted but also self-tolerant repertoire (Hogquist & Jameson, 2014). DP thymocytes passing these filters become CD4 or CD8 single-positive thymocytes that will finally leave the thymus to seed the periphery.

β-selection of DN thymocytes and subsequent positive/negative selection of DP thymocytes strongly depend upon signaling via the pre-TCR and the TCR, respectively (Cheng et al., 1997; Gascoigne, Rybakin, Acuto, & Brzostek, 2016; Molina et al., 1992; Pivniouk et al., 1998; Zhang et al., 1999). TCR signaling on DP thymocytes is activated by the receptor interaction with MHC class I or MHC class II proteins presenting self-peptides (pMHC) (Viret & Janeway, 1999). The pre-TCR instead seems to signal independently of extracellular ligands as supported by different studies. DN thymocytes expressing truncated forms of the extracellular domains of the TCRβ and/or pre-Tα chains still undergo maturation from the DN to DP stage (Irving, Alt, & Killeen, 1998; Jacobs et al., 1994), suggesting that pre-TCR signaling likely depends on the transmembrane and intracellular domains of the complex. The crystal structure of the extracellular TCRβ and pTα domains suggests a dimer of dimers with a membrane-parallel orientation that would preclude ligand binding (Pang et al., 2010). In addition, the pre-TCR internalizes at a high rate in absence of deliberate stimulation, a process that the TCR only displays upon deliberate stimulation with its pMHC ligand (Carrasco, Navarro, & Toribio, 2003; Panigada et al., 2002; Yamasaki et al., 2006). The molecular mechanisms underlying this ligand-independent signaling may involve preferential localization of the pre-TCR in signaling molecule-enriched raft domains (Saint-Ruf et al., 2000) and/or homotypic dimerization of the pre-TCR dependent on charged residues in the extracellular domain of pTα (Ishikawa, Miyake, Hara, Saito, & Yamasaki, 2010; Yamasaki et al., 2006). Still, other studies have shown that the pre-TCR can interact with MHC class I molecules in vitro (Das et al., 2016; Mallis et al., 2015), although the exact physiological implications of these interactions are unknown.

We previously reported on a Leu to Ala mutation in position 19 of the transmembrane domain of the TCR-associated CD3ζ chain (L19A) that reduces the sensitivity of effector and memory T cells for their antigenic pMHC by impairing formation of TCR nanoclusters (Kumar et al., 2011). Here, we analyze transgenic mice expressing this mutant CD3ζ chain and focus on early T cell differentiation as a model to understand the role of the pre-TCR and the mechanisms underlying its signaling capacity. Our data show that the mutation impairs pre-TCR-dependent functions essential for the early maturation of thymocytes and that the mutated CD3ζ is a hypomorphic allele with reduced intrinsic and downstream signaling capacity. This reduced signaling is associated with apparently less pre-TCR nanoclustering and strongly supports a ligand-independent mechanism for pre-TCR activation. These data further indicate an important role for the pre-TCR in shaping the repertoire of DP thymocytes, and provide a conceptual framework for understanding the emergence of autoimmune T cell repertoires.

## Results

### L19A mice have a defect in early thymic αβT cell differentiation

We generated transgenic mice expressing the previously described WT or L19A mutant murine CD3ζ-GFP fusion proteins (Kumar et al., 2011) under the control of the human CD2 promoter and enhancer, shown to direct T cell-specific expression from the earliest stages in the thymus and onwards (Greaves, Wilson, Lang, & Kioussis, 1989). Two independent founder lines were established for each transgene and backcrossed to a *Cd3z* knock-out background (Love et al., 1993), so that the transgenic CD3ζ-GFP chains were the only CD3ζ chains expressed in these mice. The mice resulting from these crosses will be referred to as WT and L19A mice throughout the text and the results shown for the WT and L19A mice combine the data obtained for the two independent WT and L19A lines, respectively, as we did not observe phenotypic differences between the two lines of each genotype (Supplementary Fig. 1a). As previously observed in mice reconstituted with CD3ζ chain transgenes under control of the human CD2 promoter and enhancer transgenic cassette, both the WT and the L19A CD3ζ chains were expressed homogeneously in all (pre-)TCR-expressing thymus populations (Supplementary Fig. 1b).

Thymi isolated from L19A mice were smaller and contained 2- to 3-fold fewer cells than thymi from WT mice (Fig 1a). The DP, 4SP and 8SP populations were reduced in number, indicating a defect in thymocyte development already occurring at the immediately previous DN stage in L19A mice (Fig 1b, left graph). In agreement, we found an increased percentage of DN thymocytes in the L19A mice as compared to WT mice (Fig 1b, right graph). Subdivision of the DN population into the DN1 - DN4 subsets using the CD44 and CD25 markers (Fig 1c) revealed similar percentages and numbers of CD44^+^CD25^-^ DN1 and CD44^+^CD25^+^ DN2 cells in L19A and WT thymuses, but a relative accumulation of CD44^-^CD25^+^ DN3 cells and a relative and absolute reduction of CD44^-^ CD25^-^ DN4 cells in L19A thymi as compared to WT thymi (Fig 1d). This showed that the earliest observable defect in αβT cell differentiation in L19A mice occurred at the transition from the DN3 to DN4 stage. Even though the L19A CD3ζ chain impaired DN3 to DN4 transition, it maintained partial functionality. The absolute numbers of DN4 thymocytes, and the percentages of DN3 and DN4 thymocytes were very low in L19A mice, but still significantly different from those of mice lacking expression of CD3ζ, which present an almost total loss of the DN4 population (Fig 1e).

**Figure 1.**
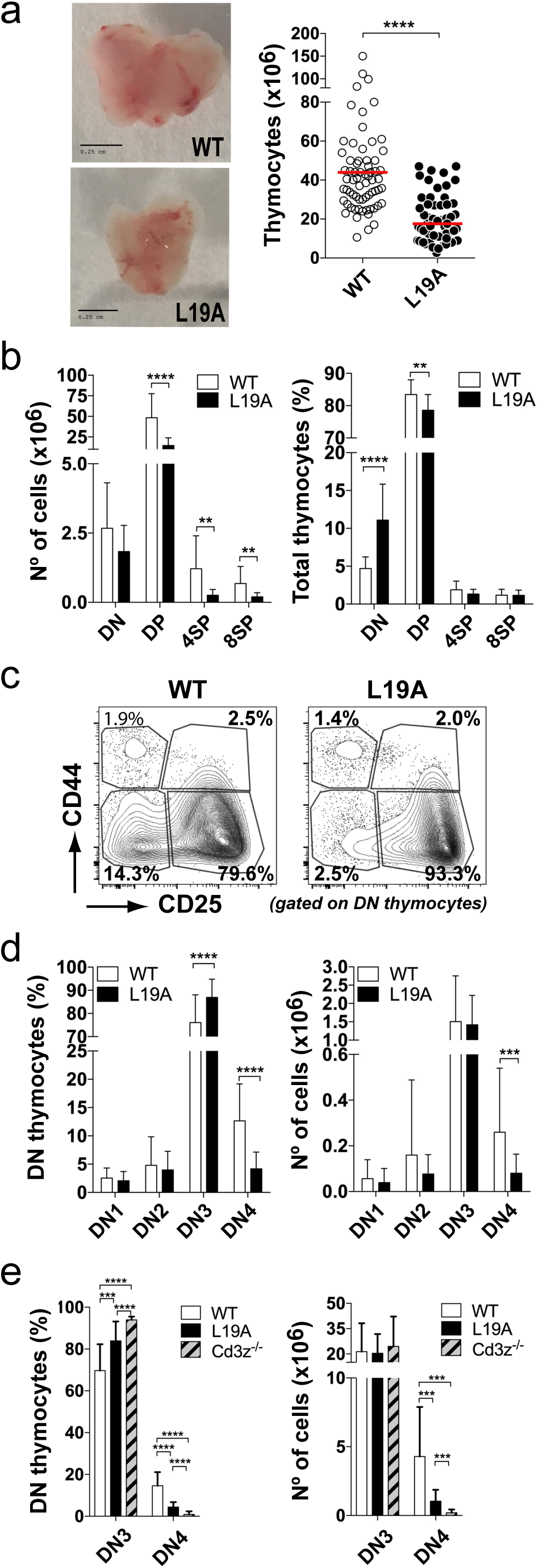
Reduction in size of thymic populations in L19A mice. (a) Thymus size and total number of thymocytes. Representative images of WT and L19A thymuses (left). Scale bar is equivalent to 0.25cm. Quantification of total number of thymocytes (right). Each dot shows the total number of thymocytes in an individual mouse. (b) Quantification of numbers (left) and percentage (right) of the major populations of thymocytes defined according to expression of CD4 and CD8. Results and statistical analysis shown are based on data from 10 independent experiments. (c) Representative plots of the DN (CD4-CD8-) subpopulations defined by the expression of CD44 and CD25 and (d) quantification in percentage (left) and cell number (right) of these populations. The DN population was pre-gated on the lineage negative (LIN-) population (CD4-, CD8-, CD19-; B220- and CD11c-). Graphs show the averages calculated on the basis of data obtained from 3 independent experiments. (e) Percentage and absolute numbers of DN3 and DN4 thymocytes in WT, L19A and *Cd3z*-deficient (Cd3z^-/-^) mice. Data come from pooling data from 38 WT, 40 L19A and 13 *Cd3z*^-/-^ mice. Graphs presents the mean ± SD. P-values were calculated using an unpaired two-tailed Student’s t test with 95% CI (* p<0.05; **p<0.01; ***p<0.001; ****p>0.0001).

### The L19A mutation impairs pre-TCR function

The DN3 to DN4 transition depends on correct formation and function of the pre-TCR. Rearrangement of the TCRβ chain occurs during the DN3 stage. DN3 thymocytes that have rearranged the TCRβ chain (DN3b thymocytes) can be distinguished from their immediate precursors (DN3a thymocytes) by a series of phenotypical changes including an increase in cell size (Hoffman et al., 1996) and upregulation of CD27 and CD28 (Taghon, Yui, Pant, Diamond, & Rothenberg, 2006; Williams et al., 2005). Applying this first criterion, we quantified the percentage of DN3a and DN3b thymocytes in the DN population of L19A and WT mice using as a reference DN3 thymocytes from *Cd3e*-deficient mice which, even though they can rearrange their *Tcrb*-locus, cannot execute the pre-TCR-dependent differentiation steps and are thus virtually unable to reach the DN3b stage (DeJarnette et al., 1998). L19A thymuses contained significantly more DN3a and fewer DN3b thymocytes than WT thymuses. However, this block in differentiation was not as severe as the one observed in *Cd3e*-deficient thymuses, suggesting inefficient differentiation rather than a full blockade (Fig. 2a). This lag in differentiation was not due to the inability of L19A thymocytes to rearrange their *Tcrb*-locus, as intracellular staining for TCRβ in the DN3b and DN4 populations of WT and L19A thymuses showed an equal intensity (Fig. 2b). Moreover, cell surface expression of the pre-TCR, measured by extracellular staining with antibodies against the TCRβ and pTα chains, on the DN3b-enriched CD44^-^CD25^lo^ population and CD44^-^CD25^-^ DN4 cells, was indistinguishable between L19A and WT mice (Fig 2c and d). Likewise, cell surface expression of CD3ε and expression of the transgenic CD3ζ-GFP chain was equal for WT and L19 DN3b and DN4 thymocytes (Supplementary Fig. S2). Thus, L19 mutant thymocytes demonstrated no defects in formation and cell surface expression of the pre-TCR but showed reduced numbers of cells successfully transiting to the DN4 compartment.

**Figure 2.**
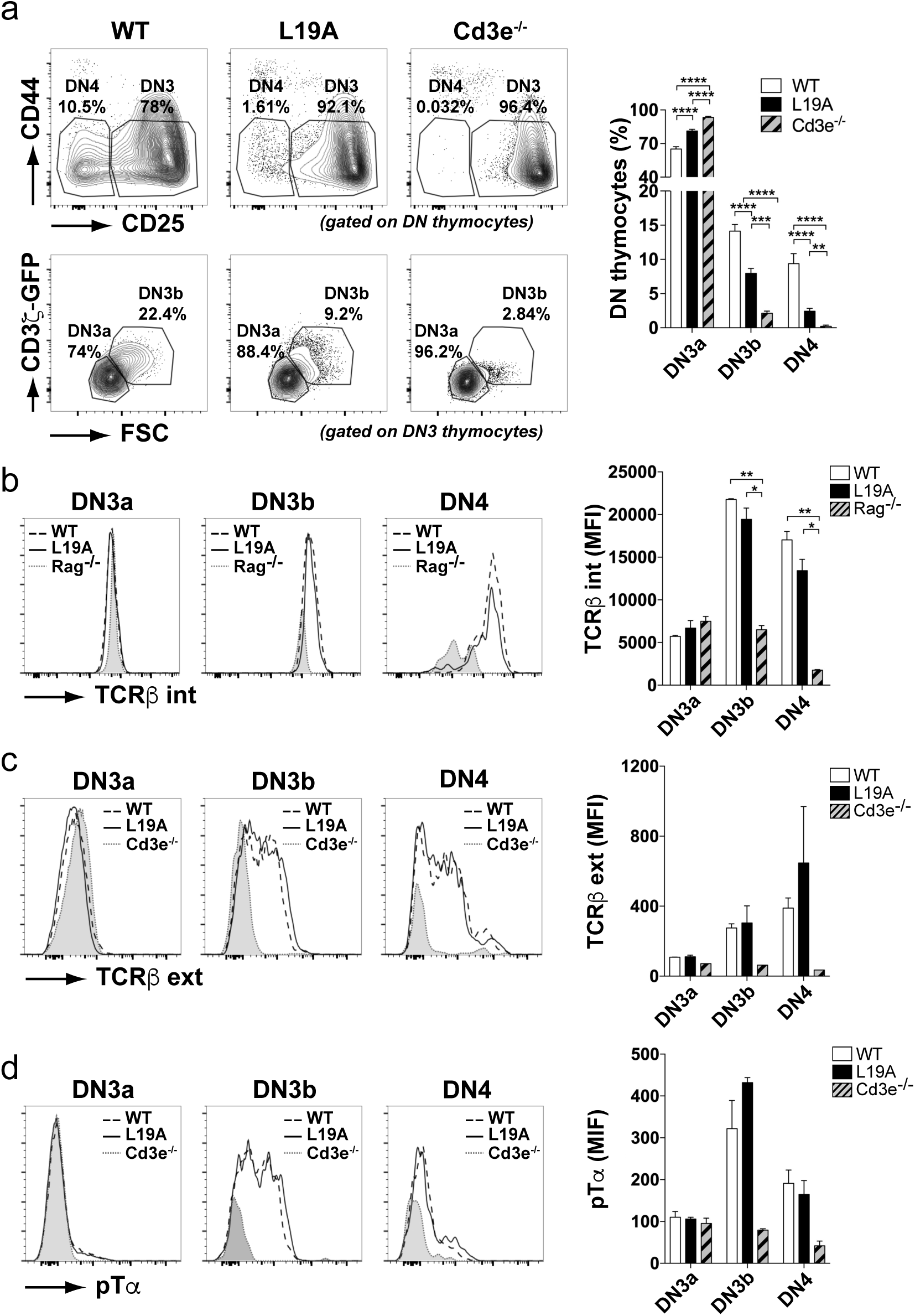
T cell differentiation in L19A mice is impaired at the DN3a to DN3b transition. (a) Representative flow cytometry plots of double negative (DN) populations in WT, L19A and *Cd3e*-deficient (Cd3e^-/-^) thymuses according to CD44 and CD25 markers (upper panels; CD25^+/lo^CD44^-^ DN3 and CD25^-^CD44^-^ DN4 gates indicated) and identification of DN3a and DN3b subpopulations within the DN3 gate according to forward scatter (FSC) and CD3ζ-GFP expression (bottom panels). On the right, quantification based on pooled data from six independent experiments showing the percentage of the DN3a, DN3b and DN4 populations. (b) Representative histograms (left) and quantification (right) of the intracellular TCRβ expression levels in the DN3a (CD44^-^CD25^+^), DN3b (CD44^-^CD25^lo^) and DN4 (CD44^-^CD25^-^) subpopulations. Thymocytes from *Rag1*-deficient mice were used as a control for intracellular staining and quantification is based on one out of four experiments. (c) Representative histograms (left) and quantification (right) of the extracellular TCRβ (top panels). Data in graph represent one out of four experiments. (d) pTα expression level for the indicated DN subsets, defined by level of CD25 expression. Data in graphs are based on one out of 3 experiments. Graphs present mean ± SEM. P-values were calculated using an unpaired two-tailed Student’s t test with 95% CI (* p<0.05; **p<0.01; ***p<0.001; ****p>0.0001).

Expression of the pre-TCR at the cell surface promotes survival, proliferation and differentiation to the DP stage (von Boehmer, 2005). We measured these processes in WT and L19A DN thymocytes and used thymocytes from *Cd3e*^*-/-*^ mice as a control. Upregulation of the transferrin receptor (CD71) and the large neutral amino acid transporter subunit CD98 was significantly impaired in DN3b and particularly in DN4 thymocytes of L19A mice (Fig 3a). In L19A mutants, BrdU-positive, proliferating DN3 and DN4 thymocyte were clearly diminished in numbers as compared to the corresponding WT thymocytes (Fig 3b) and they were also more prone to undergo apoptosis (Fig 3c). Together, these results clearly indicated that the function of the pre-TCR in L19A mice was impaired.

**Figure 3.**
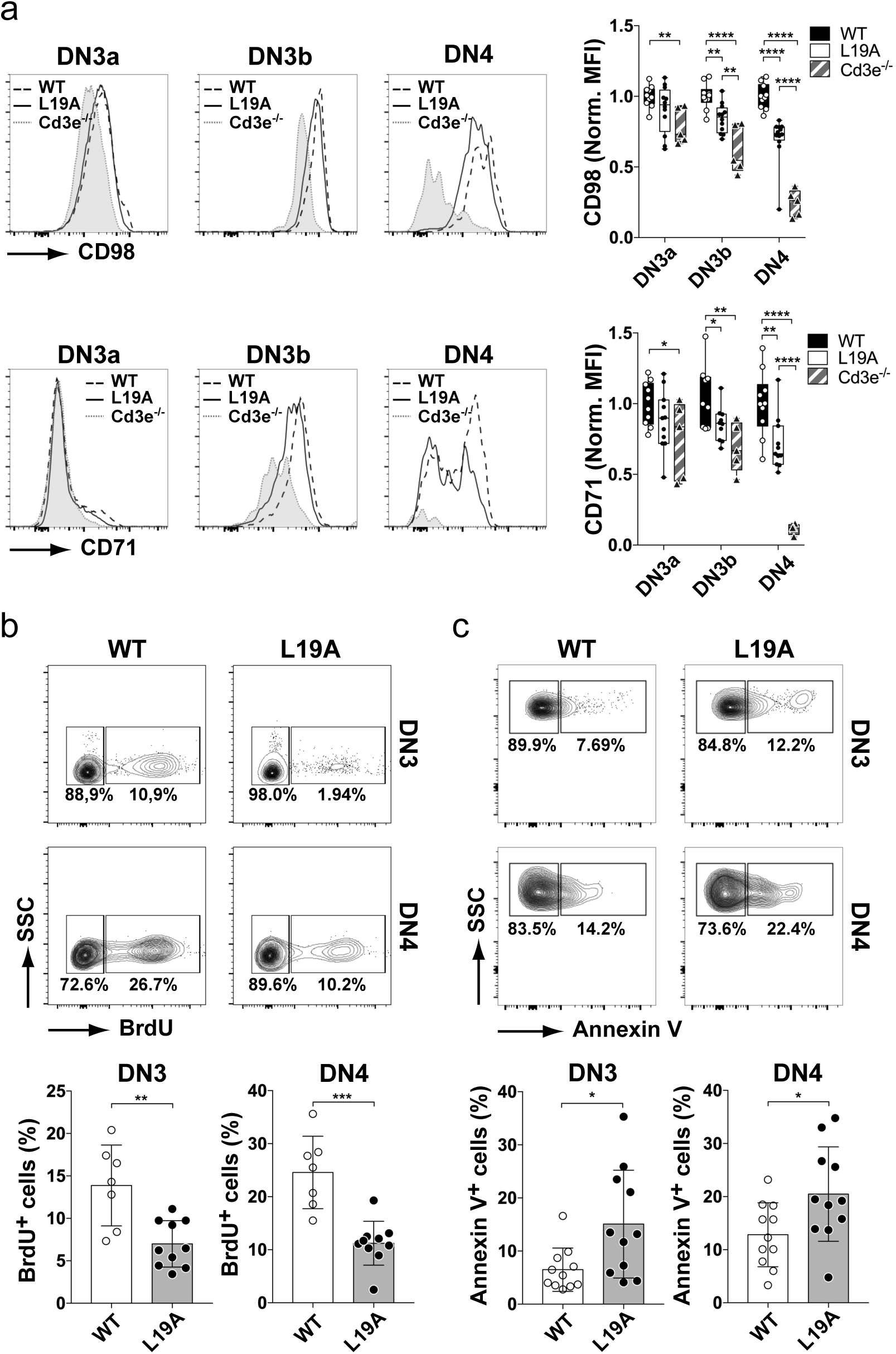
The CD3ζ transmembrane region regulates pre-TCR-dependent thymocyte differentiation and proliferation. (a) Representative plot and quantification of expression of the CD98 (top panels) and CD71 (bottom panels) maturation markers at the cell surface of DN3 and DN4 thymocytes. CD44^-^ DN subsets were gated according CD25 levels: DN3a (CD25^lo^), DN3b (CD25^lo^) and DN4 (CD25^-^). Graphs present quantification of pooled data from four independent experiments. (b) Representatives plots of BrdU staining in DN3 and DN4 populations and quantification of the percentage of BrdU+ thymocytes within DN3 (left) and DN4 (right) populations. Thymocytes shown in the plots were pre-gated on Lin-CD44- population. Graphs present the mean ± SEM of pooled data from three independent experiments. (c) Representative plots and quantification of Annexin V binding in DN3 and DN4 populations. Thymocytes shown in the plots were pre-gated on the Lin-CD44- population. Quantification shows the mean ± SEM of pooled data from four independent experiments. P-values were calculated using an unpaired two-tailed Student’s t test with 95% CI (* p<0.05; **p<0.01; ***p<0.001; ****p>0.0001).

### The CD3ζ transmembrane domain controls pre-TCR mediated signaling efficiency

To understand the mechanisms underlying the compromised pre-TCR function in L19A thymocytes, we first compared their pre-TCR signaling capacity with that of WT thymocytes. We determined the levels of activation of different signaling components directly *ex vivo*, without deliberate stimulation with anti-CD3 or -TCRβ antibodies, in order to closely resemble the physiological signaling status (Fig. 4a). To this end we first focused on a proline-rich sequence (PRS) of the intracellular domain of CD3ε. This motif is selectively exposed in active TCRs and can be detected via intracellular staining with the APA1/1 mAb (Blanco, Borroto, Schamel, Pereira, & Alarcon, 2014; Risueno, Gil, Fernandez, Sanchez-Madrid, & Alarcon, 2005). We used DN thymocytes isolated from KI-PRS mice as a negative control for staining, as the key proline residues in the CD3ε PRS of these mice are mutated to alanines, abrogating recognition by the APA1/1 mAb (Borroto et al., 2013). Freshly isolated DN3b and DN4 thymocytes from WT and L19A mice, identified as CD44^-^CD25^lo^ and CD44^-^CD25^-^ DN thymocytes respectively, showed above-background staining, indicating actively signaling pre-TCRs (Fig. 4b). However, L19A DN3b and DN4 thymocytes showed a significantly lower APA1/1 staining intensity than WT DN3b and DN4 thymocytes. This was not due to a lower level of expression of CD3ε in L19A thymocytes (see Supplementary Fig. 2), and hence indicated fewer active pre-TCRs in L19A thymocytes. On the other hand, we observed a high and equal level of APA1/1 staining in WT and L19A CD44^-^CD25^hi^ DN3a thymocytes. DN3a thymocytes have not yet rearranged the TCRβ chain but do express CD3ε (see Immune Genome website: https://www.immgen.org/; (Mingueneau et al., 2013)). In absence of TCRβ the CD3ε chains are retained in the ER and are in the conformation recognized by APA1/1 (Borroto, Mallabiabarrena, Albar, Martinez, & Alarcon, 1998). Thus, equal levels of APA1/1 staining indicate cytoplasmic retention of CD3ε chains in L19A mutant DN3 thymocytes.

**Figure 4.**
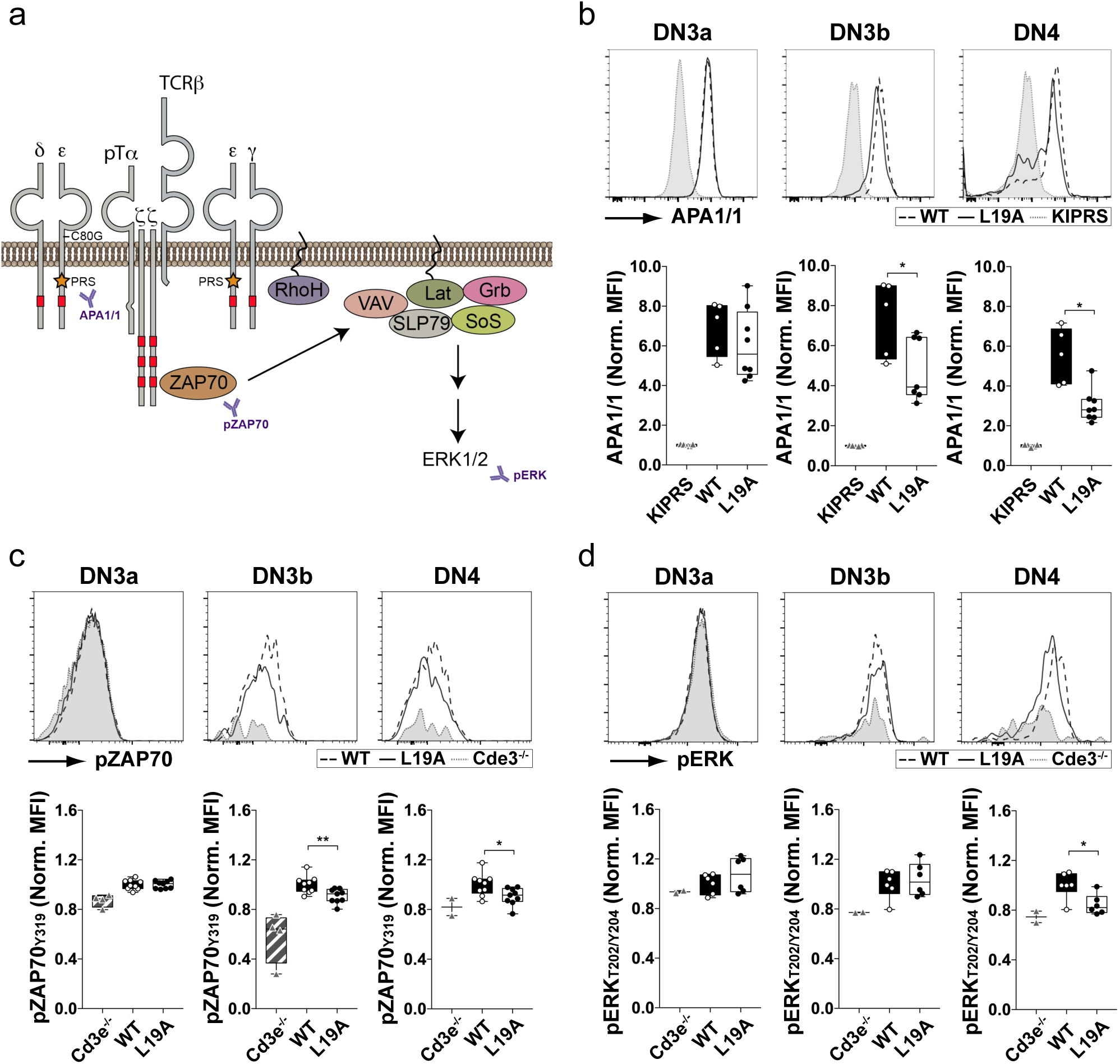
Pre-TCR signaling by L19A thymocytes is reduced compared with WT counterparts. (a) Scheme of pre-TCR signaling pathways. (b) Representative histograms and normalized quantification of intracellular APA1/1 staining. Graphs show the mean ± SEM of pooled data from two independent experiments. (c) Representative histograms and normalized quantification of intracellular pZAP70 (P-Y319) labeling. Graphs show the mean ± SEM of three independent experiments. (c) Representative histograms and normalized quantification of pERK (P-Thr202/P-Tyr204) staining. Data show the mean ± SEM of two independent experiments. P-values were calculated using an unpaired two-tailed Student’s t test with 95% CI (* p<0.05; **p<0.01).

We also measured phosphorylation of the receptor-proximal signaling molecule ZAP70 and of the MAP kinase ERK downstream of the pre-TCR. As for the PRS exposure measurements, phosphoprotein-specific labeling was performed on freshly isolated thymocytes and the DN3a, DN3b and DN4 were identified with the CD25 and CD44 markers. Thymocytes from mice lacking the CD3ε chain were used as a background control. L19A DN3b and DN4 thymocytes showed decreased phosphorylation of Y319 of ZAP70 as compared to WT thymocytes (Fig. 4c). In DN3a thymocytes no such differences were found. The staining intensity was undistinguishable between L19A mutant, WT, and CD3e^-/-^ thymocytes (Fig. 4b), as DN3a thymocytes from all three genotypes lack a functional pre-TCR. Finally, L19A DN4 thymocytes demonstrated impaired Thr202/Tyr204 phosphorylation of ERK signaling (Fig. 4d) as compared to WT thymocytes. Altogether, our data indicate reduced activation levels within and downstream of the pre-TCR in L19A thymocytes compared to the WT ones, thus showing a role for pre-TCR signaling intensity in the quantitative output of the β-selection process.

### Receptor confinement and diffusion

We next analyzed pre-TCR diffusion and confinement using TIRF microscopy. Comparison of the dynamics of pre-TCR at the cell membrane provides indications of its function and signaling capacity. DN3 and DN4 thymocytes were purified in two sequential steps, first via negative selection of DN thymocytes by magnetic depletion of DP and lineage-positive (Lin^+^) thymocytes, followed by sorting for CD44^-^ DN thymocytes with a maximal threshold of GFP expression (Fig. 5a). Selected thymocytes were adhered to fibronectin-coated cover slips and analyzed in a Leica AM TIRF inverted microscope in TIRF mode. We used the CD3ζ-coupled GFP as the label to track the pre-TCR complexes. Analysis of cell surface labeling of WT and L19A DN thymocytes showed a strong correlation between GFP and CD3ε staining (Supplementary Fig. 2d), validating the use of the GFP label to track the pre-TCR. Only particles that were continuously detectable and did not fuse or split during a time period of at least 5 seconds were analyzed. Data showed an increased diffusion rate of the L19A pre-TCR particles as compared to WT particles (Fig. 5b). When analyzing the tracks of the mobile particles for confined, free and directed diffusion no significant differences in distribution between these modes of diffusion were observed between surface particles of WT and L19A DN thymocytes (Fig. 5c). This indicated that the overall speedup was unlikely due to a mutation-dependent shift in the mode of diffusion across the membrane. Notably, the increased diffusion rate of the L19A particles was accompanied by decreased fluorescence intensity, a measure that correlates with the number of receptors within each fluorescent particle (Fig. 5d). In addition, L19A thymocytes had a reduced fraction of immobile pre-TCR particles as compared to WT DN thymocytes (Fig. 5e), suggesting that the mutation also interfered with the localization of pre-TCRs in certain membrane domains or impeded association with membrane-associated structures. Together these data suggest that WT pre-TCR aggregate to form pre-TCR nanoclusters and that the presence of L19A reduces both the mobility and fluorescent intensity of these nanoclusters.

**Figure 5.**
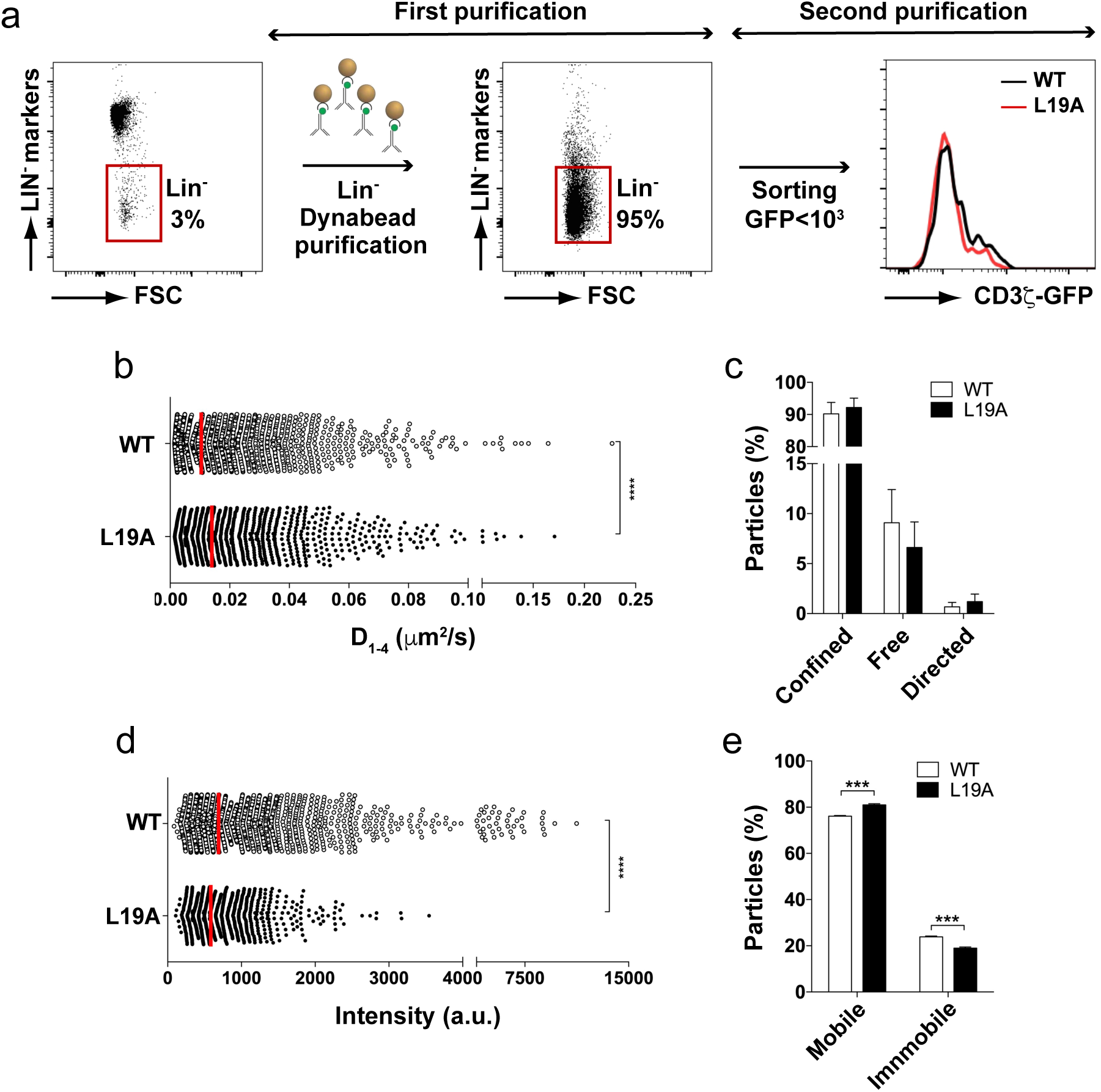
TIFRM analysis of the pre-TCR on CD44^-^ primary thymocytes. (a) Purification strategy of DN CD44^-^ thymocytes through two purification steps, depleting Lin^+^ thymocytes from pools of three thymi of each transgenic mouse line, followed by sorting CD44^-^ LIN^-^ cells with a cut-off of GFP fluorescence intensity of 10^3^ relative units. (b) Median of the short time-lag diffusion coefficient (D_1-4_) of the mobile particles detected in the analysis. (c) Quantification of the percentage (mean ± SEM) of mobile particles for confined, free and directed diffusion tracks (d) Quantification of the mean fluorescent intensity of each spot during the first 20 frames in which the particles are visible. Each dot in the graph presents single CD3ζ-GFP spot (e) Percentage (mean ± SEM) of mobile and immobile particles in the WT and L19A samples. Pooled data were obtained in 4 independent experiments encompassing on average 19.1 and 13.1 tracked particles per cell for 131 WT cells (range 20 - 57 cells/experiment) and 110 L19A cells (range 13 - 40 cells /experiment), respectively. P-values were calculated using an unpaired two-tailed Student’s t test with 95% CI (***p<0.001; ****p>0.0001).

### TCRβ repertoire diversity

The impairment in signaling observed for the L19A pre-TCR might not only give rise to reduced thymic population sizes but also alter the DP pre-selection repertoire. We purified large CD69^-^ DP thymocytes, that have not yet been subject to negative and positive selection from WT and L19A mice and obtained on average 1.4 ± 0.8 × 10^5^ WT and 0.4 ± 0.3 × 10^5^ L19A cells (mean ± SD; n=5 each; p=0.032 Mann-Whitney) (Fig. 6a). We determined the V, D and J segments used by each TCRβ chain via an unbiased 5’ RACE protocol and identified its CDR3 region as described in material and methods. An average of 8.4 ± 2.0 × 10^5^ and 4.3 ± 1.4 × 10^5^ (mean ± SD; n=5 each; p=0.016, Mann-Whitney) of in-frame VDJ reads were obtained for the WT and L19A libraries, respectively. In order to estimate the number of clones represented by these reads, we took into account the error rate of the Illumina sequencing methodology (Shendure & Ji, 2008) and applied a filter to each library, grouping reads that differed in less than 3% of their CDR3 nucleotide sequence. These data sets were used for all subsequent analyses of the diversity of the TCRβ repertoire. We determined the effective number of clones in each repertoire by calculating Hill numbers ^*q*^*D* via rarefaction and extrapolation methods (Chao et al., 2014), being ^*0*^*D* the “Richness” or effective number of different clones. For other values of q, ^q^D estimates the diversity when giving more weight to more frequent clones (large positive q) or rare ones (negative q). Rarefaction curves for the WT DP repertoires separated almost completely from the curves for L19A DP repertoires, indicating that repertoires were more diverse in WT mice than in L19A mice (Fig. 6b). The observed number of clones (or richness diversity, ^*0*^*D*) was 7.6 ± 4.5×10^4^ for WT and 2.3 ± 1.5×10^4^ clones for L19A (mean ± SD; p=0.032, Mann-Whitney) and the estimated richness was 7.9 ± 4.7×10^4^ for WT and 2.4 ± 1.5×10^4^ (mean ± SD; p=0.031, Mann-Whitney). When calculating diversity using q values of 1 and 2, thereby providing more weight to the more abundant clones within the populations, we found an essentially identical pattern of diversities (Supplementary Fig. 3a and b). The latter provided evidence for the robustness of the analysis of early DP repertoire diversity. The coverage (Chao et al., 2014) for each mouse was above 0.98 so the extrapolated parts of the curve were close to the asymptotic diversity and guaranteed that the estimation of the diversity of the repertoires was accurate (Supplementary Fig. 3c).

**Figure 6.**
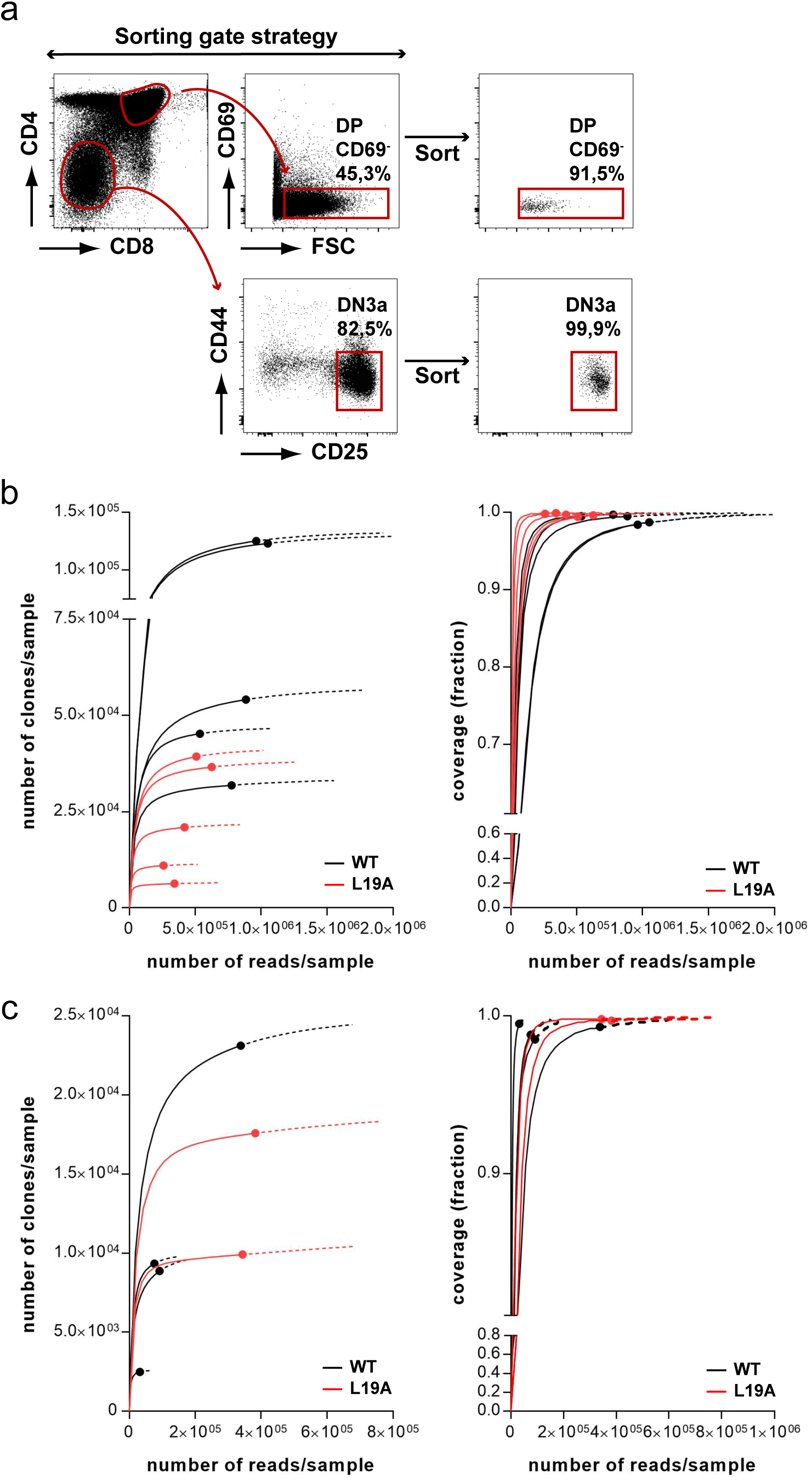
TCRβ diversity of DP and DN thymocytes in WT and L19A mice. (a) Sorting strategy for isolation of early DP and DN3 thymocytes. (b) Number of clones per sample estimated with the richness diversity (^*0*^*D*) for DP thymocytes. Solid lines: Rarefaction (interpolation) curve; dashed lines: extrapolation curves; symbols: observed diversity. Note that the observed values are, in all cases, close to the asymptotic part of the curves, consistent with the large values of the coverage (fig. S3). (c) Number of clones per sample estimated for DN3 thymocytes. Meaning of lines and symbols as in (b).

We also compared diversity of the TCRβ repertoire in WT and L19A CD25^hi^CD44^-^ DN3 thymocytes that are enriched for cells that have rearranged the *Tcrb* gene segments but should have undergone little pre-TCR mediated selection and expansion (Fig. 6a). We obtained TCRβ sequences from 4 WT and 2 L19A samples containing 8.6 ± 5.2 × 10^4^ and 8.2 ± 1.9 × 10^4^ cells (mean ± SD; p=0.800 Mann-Whitney) and giving rise to 1.3 ± 1.4 × 10^5^ and 3.6 ± 0.3 × 10^5^ filtered, in-frame reads (mean ± SD; p=0.133 Mann-Whitney) for WT and L19A samples, respectively. No clear separation of rarefaction curves derived from the WT and L19A DN3 repertoires was observed, supporting the notion that diversity of these repertoires was similar (Fig. 6c). Indeed, the observed number of clones in WT and L19A DN repertoires was not significantly different (1.1 ± 0.9×10^4^ and 1.4 ± 0.5×10^4^ for L19A clones for WT and L19A repertoires, respectively; mean ± SD; p=0.53 Mann-Whitney). We could not detect differences in CDR3 length or V, D and J usage accompanying the difference in diversity between WT and L19A early DP thymocytes (Supplementary Fig. 3d and e), indicating that the nature of the TCRβ chain did not influence the ability of a DN3 thymocyte to become an early DP thymocyte. Together these data show that the intensity of signaling via the pre-TCR influences the diversity of the TCRβ repertoire that can undergo subsequent positive and negative selection.

## Discussion

The intensity of signaling via the TCR determines the fate of DP thymocytes during positive and negative selection, thereby assuring the generation of a repertoire of mature T cells that is MHC-restricted and self-tolerant (Hogquist & Jameson, 2014). The studies employed to reach these conclusions have been greatly facilitated by the availability of pMHC ligands with known affinities for defined TCRs. It has been much more difficult to establish whether pre-TCR signaling intensity plays a role in the transition from the DN3 to the DN4 and early DP stages and whether this has consequences for the repertoire of thymocytes that can undergo positive and negative selection. The inexistence or lack of defined ligands and the small number of DN3 and DN4 cells present in the thymus have hampered the study of pre-TCR signaling in primary thymocytes. The relevance of this study with regard to understanding the role of pre-TCR signaling intensity is thus three-fold. We measured pre-TCR signaling in primary DN3 and DN4 thymocytes directly after isolation from the thymus and without deliberately stimulating the pre-TCR, thereby obtaining data sets that closely reflect the *in vivo* conditions. We demonstrated reduced activation levels of defined signaling components within and downstream of the pre-TCR in L19A thymocytes, as compared to WT thymocytes, and therefore show a role for pre-TCR signaling intensity in the quantitative output of the β-selection process. We lastly provided direct evidence that the intensity of pre-TCR signaling has a quantitative impact on the TCRβ diversity of the repertoire of early DP thymocytes that still need to undergo positive and negative selection. This latter finding implies that during β-selection not all DN precursors with in-frame rearranged *Tcrb* genes mature but that pre-TCR-signaling efficiency imposes a filter on the emerging repertoire. The data on the dynamic behavior of WT and L19A pre-TCRs, together with our previous findings on the L19A mutation in context of the TCR, shed light on the molecular mechanisms underlying pre-TCR signaling.

We identify the transmembrane domain of CD3ζ as a regulator of pre-TCR function. Given the nature of this domain and taking into account our previous data, we envision that CD3ζ can affect pre-TCR function in two not mutually exclusive ways, namely nanocluster formation and guidance of the pre-TCR towards lipid rafts. TCR nanoclusters, groupings of up to 20 TCRs present at the cell surface of T cells before interaction with its pMHC ligands, have been detected via biochemical, high resolution fluorescence microscopy and immuno-EM approaches by different research groups (Lillemeier et al., 2010; Schamel et al., 2005; Sherman et al., 2011; Zhong et al., 2009), although their existence has been disputed (Brameshuber et al., 2018; Rossboth et al., 2018). The L19A mutation leads to a reduced formation of TCR nanoclusters in mature T cells. T cells expressing this mutant CD3ζ chain have reduced sensitivity for their cognate pMHC (Kumar et al., 2011). CD3ζ could also allow formation of pre-TCR nanoclusters. Our findings using single particle tracking in TIRFM are compatible with the existence of such nanoclusters. The fluorescence intensity of the particles we detect is stable over the time of observation, and these particles do not split or fuse with other particles, supporting the idea that the pre-TCRs form stable clusters instead of brief dynamic accumulations. We observe particles with a range of fluorescence intensities that would be compatible with clusters of various sizes. The higher intensity of GFP-coupled CD3ζ particles observed at the cell surface of WT DN thymocytes as compared to L19A DN thymocytes indicate that the L19A mutation also reduces the size of such pre-TCR nanoclusters. The increased diffusion rate of these particles in the L19A DN thymocytes is in this case also to be expected. However, the limited resolution of TIRFM does not allow us to distinguish between the possibility that the observed particles represent pre-TCR nanoclusters of various sizes or whether these are stable membrane domains that contain more or less pre-TCRs. We nevertheless favor the idea that pre-TCR nanoclusters do indeed form, and thus the differentiation defect associated with the L19A mutation underscores the relevance of nanoclusters in pre-TCR signaling. Nanoclustering of the TCR can enhance TCR signaling by facilitating multivalent ligand binding (Fahmy, Bieler, Edidin, & Schneck, 2001; Martin-Blanco et al., 2018; Molnar et al., 2012) and by permitting cooperative signaling between engaged and non-engaged TCRs (Martinez-Martin et al., 2009). Notably, nanoclustering also favors one of the early key events of TCR signaling, induction of a conformational change of the intracellular domain of CD3ε (Minguet, Swamy, Alarcon, Luescher, & Schamel, 2007). This conformational change exposes the intracellular proline rich sequence (PRS) of CD3ε (Gil, Schamel, Montoya, Sanchez-Madrid, & Alarcon, 2002) and might be involved in dislodging the CD3ε intracellular domain from the inner membrane leaflet, facilitating its recognition by the TCR-associated signaling machinery (Gagnon, Schubert, Gordo, Chu, & Wucherpfennig, 2012; Xu et al., 2008). The conformational change appears critical in the case of pre-TCR-dependent signaling as β-selection is severely impaired in mice with mutations in the extracellular stalk domain of CD3ε that prevent this conformational change (Blanco et al., 2014; Y. Wang et al., 2009) and in mice with a mutated or deleted proline rich sequence (PRS) (Borroto et al., 2013; Brodeur, Li, da Silva Martins, Larose, & Dave, 2009). This is supported by our finding that the impaired transition from DN to DP thymocytes in L19A mice coincides with a reduced exposure of the PRS. Notably, the antibodies used to rescue the DN to DP transition in RAG-deficient mice (Levelt, Mombaerts, Iglesias, Tonegawa, & Eichmann, 1993) are known to induce and/or stabilize the conformational change of the CD3ε intracellular domain (Gil et al., 2002). The pre-TCR does not appear to depend on binding of external ligands to signal and therefore, induction of the conformational change would have to be regulated in a different manner. Assuming that the pre-TCR oscillates between a signaling-competent, CD3ε-exposing stage and a non-competent stage, as has been proposed for the TCR (Martin-Blanco et al., 2018)(Martin-Blanco et al), we speculate that this exposure is stabilized by its interaction with neighboring pre-TCR complexes. This interaction could in part rely on the inherent ability of the pTα extracellular domain to form homodimers, but would also need the transmembrane domain of CD3ζ, by allowing formation of pre-TCR nanoclusters.

The transmembrane domain of CD3ζ could also be involved in guiding the pre-TCR towards cholesterol-enriched membrane domains (Saint-Ruf et al., 2000). Direct evidence obtained in mature T cells shows that TCRβ expressed at the cell surface membrane can bind to cholesterol (Molnar et al., 2012). The transmembrane region of CD3ζ contains CARC and CRAC motifs that have been implied in the binding of membrane-embedded cholesterol (Fantini, Di Scala, Baier, & Barrantes, 2016). The L19A mutation is located between the amino acids defining the CRAC motif, possibly altering its ability to interact with cholesterol. The typical punctuate distribution of the pre-TCR on the cell surface membrane of the DN thymocyte-derived cell line SCB.29 and the integrity of TCR nanoclusters on the cell surface of mature T cells are sensitive to treatment of these cells with the cholesterol-extracting compound methyl-β-cyclodextrin (Saint-Ruf et al., 2000; Schamel et al., 2005). Furthermore, displacement of cholesterol by cholesterol-sulfate reduces TCR nanocluster formation and sensitivity of DP thymocytes for cognate pMHC and alters the extent of positive selection (F. Wang, Beck-Garcia, Zorzin, Schamel, & Davis, 2016). Together these data indicate the relevance of cholesterol in maintenance of these molecular assemblies and their function, and point out a potential role for CD3ζ in this process.

The degree of diversity of the repertoire of TCRβ chains is correlated with the signaling capacity of the pre-TCR, suggesting that signal quantity imposes a selective pressure on the formation of the TCRβ repertoire. This selection step does not appear to depend on the molecular features of the TCRβ chain. We did not find changes in CDR3 length between DN3 and early DP thymocytes, or between WT and L19A DN3 and early DP thymocytes, or over- or underrepresentation of particular V, D and J elements or combinations thereof, features that, if observed, could have hinted at the involvement of a ligand. This finding is in line with the paradigm that the pre-TCR functions without extracellular ligands. Pairing of the pTα/TCRβ heterodimer is mediated by contacts between the constant domain of TCRβ and the invariable pTα chain (Pang et al., 2010) and no influence of the nature of the TCRβ chain is thus to be expected from this developmentally critical step. The formation of dimers of pre-TCR heterodimers, should not be influenced either by the type of TCRβ chain expressed by the thymocyte, as this interaction depends on strongly conserved residues in the variable domain of the TCRβ chain of one heterodimer with the constant pTα chain of the other heterodimer. From our work on TCR nanoclusters we have not found evidence for differences in the ability of particular TCRβ chains to promote TCR nanocluster formation. We therefore envision the existence of a minimal threshold of pre-TCR signaling intensity that permits transition from the DN to DP stage. The ability to reach this threshold would in part be determined by the degree of nanoclustering of the pre-TCR. The L19A mutation, interfering with pre-TCR nanoclustering and thus signaling intensity, would allow fewer clones to reach the threshold as compared to a WT CD3ζ chain and give rise to a narrower repertoire of early DP thymocytes in L19A as compared to WT mice. Independent of whether this threshold exists, it is clear from our data that β-selection permits maturation of DN thymocytes with a range of pre-TCR-dependent signaling capabilities, as pre-TCRs of L19A DN3b and DN4 thymocytes show lower activation levels of intrinsic and downstream signaling components than those of WT DN3b and DN4 thymocytes.

A relevant question concerns the potential consequence of the permissiveness of differentiation of DN clones with reduced signaling capacity for the subsequent selection steps and the mature T cell repertoire. In the context of the L19A mutation it would be expected that, if clustering of the TCR of DP thymocytes occurs, the reduced signaling capacity would be maintained in these precursors and TCRs with higher self-pMHC affinity would be allowed to mature, giving rise to a potentially auto-reactive T cell repertoire. In a general context, mutation or allelic variation of components involved in signaling via the pre-TCR and/or variation in their expression level, could be expected to have similar effects on pre-TCR signaling capacity of DN thymocytes, and promote the selection of higher affinity clones upon DP differentiation. Thus, our finding that β-selection allows differentiation of clones with a range of pre-TCR signaling intensities may facilitate emergence of a self-biased repertoire. With regard to the effects of the lower diversity of TCRβ clones, this could result in a lower diversity of the mature αβTCR repertoire, although it is not straightforward to estimate. Positive and negative selection impose a strong limit on the number of DP clones that differentiate into mature T cells (Merkenschlager et al., 1997; van Meerwijk et al., 1997) and a thorough quantitative analysis of the diversity of the mature T cell repertoire would be needed to establish the relative contribution of positive/negative selection and the here-described pre-TCR limiting effect on repertoire diversity.

Altogether, our finding that the diversity of the repertoire of TCRβ sequences is reduced in mice with impaired pre-TCR signaling points to an additional selective pressure on DN negative thymocytes during β-selection, potentially involving selection of thymocytes that have a configuration of the pre-TCR and downstream signaling machinery that allows them to reach a signaling threshold necessary for progressing to the subsequent stages of differentiation. At the same time our data show that DN thymocytes with a range of signaling capabilities can progress in their differentiation, which, we speculate, give rise to repertoires with more or less self-recognizing tendencies. In this regard, the organization of the pre-TCR at the cell surface plays a critical role.

## Materials and Methods

### Mice

CD3ζWT-GFP and CD3ζL19A-GFP transgenic mouse lines were generated in the CNB/CBMSO transgenic facility. The previously described CD3ζWT-GFP and CD3ζL19A-GFP cDNA sequences (Kumar et al., 2011) were amplified with 5’ and 3’ flanking oligos containing an *Eco*RI and *Sal*I site, respectively, and cloned into the *Eco*RI and *Sal*I sites of the transgenic cassette vector p29Δ2 in which expression of the inserted DNA fragment is regulated by the hCD2 promoter and enhancer (Greaves et al., 1989). Transgenic mice were obtained by injecting the promoter, transgene and enhancer sequences-containing *Not*I fragment into C57BL/6JxSJL F1 1-cell stage embryos and transplanting the embryos in pseudo-pregnant foster mothers. Two male founder mice for each transgenic construct (WT-F10, WT-F19, L19A-F13 and L19A-F23), initially identified via southern blot using a GFP-specific probe, were selected and backcrossed once with C57BL/6J females to check for germ line transmission and further backcrossed to a *Cd3z* knock-out background (Love et al., 1993). Relative copy numbers of the transgenes, as determined by quantitative PCR analysis of genomic DNA, were 3-5 fold higher for the L19A lines as compared to the WT lines, which displayed similar copy numbers amongst themselves (fig S1c). Mice backcrossed for 2 to 5 generations to the C57BL/6J background, homozygous for the *Cd3z*^*tm1Lov*^ allele and always carrying the transgene in hemizygosity were used for the experiments described in this manuscript. Knock-in mice with a PxxP to AxxA double mutation in the PRS of CD3ε were described previously (Borroto et al., 2013). *Cd3e* knock-out mice were obtained from Jackson Laboratories (stock #004177) (DeJarnette et al., 1998). Mice homozygous for the *Rag1*^*tm1Mom*^ knock-out allele (Mombaerts et al., 1992) were kindly provided by Dr. César Cobaleda (CBMSO, Madrid). Mice were maintained under SPF conditions in the animal facility of the CBMSO in accordance with applicable national and European guidelines. All animal procedures were approved by the ethical committees of the Consejo Superior de Investigaciones Científicas and the Comunidad Autónoma de Madrid (PRO-EX 68/14 and PRO-EX 384/15).

### Antibodies and reagents

Antibodies used to detect the following mouse proteins were: CD4-PerCP, -647, -FITC (RM4-5), CD8-biotin, -PerCP, -647 (53-6.7), CD11b-biotin (M1/70), CD16/32 purified (2.4G2), CD19-biotin (1D3), CD25-PerCP, -APC, -PE (3C7), Gr1-biotin (RB6-8C5), CD44-BV421, -PE and -APC (IM7); TCRβ-APC, -PE, and -BV421 (H57-597); CD3ε-PerCP, -APC and -PE -biotin (2C11), NK1.1-biotin (PK136), pTα purified (2F5), CD3γε-APC (17A2), anti-mouse pZAP70 -647 (Y319), obtained from BD Pharmingen; F4/80-biotin (BM8), CD98-PE (RL388), CD8-BV421 (53-6.7), CD71-PE (R71217), all from eBioscience. Annexin V-PE, 7AAD and the APC-labeled anti-BrdU mAb (3D4) were purchased from BD Pharmingen. The APA1/1 monoclonal antibody, which recognizes a conformational epitope of CD3ε, and its use as a probe for the conformational change of the TCR have been described previously (Risueno et al., 2005)(Risueno et al., 2005). Where necessary, secondary antibodies (anti-rabbit Alexa647 and anti-mouse Alexa647 from ThermoFisher) or fluorescent probes (Streptavidin-Percp and -APC from BD Pharmingen, Streptavidin-PE from Invitrogen and Streptavidin -BV421 from Biolegend) were used.

### Flow cytometry

Thymi from 5-9 week-old mice were homogenized with 40 μm strainers and washed in phosphate-buffered saline (PBS) containing 1% bovine serum albumine (BSA). Single-cell suspensions were counted with a Scepter 2.0 Cell Counter and incubated with fluorescence-labelled antibodies for 30 minutes at 4°C after blocking Fc receptors using an anti-CD16/32 antibody. For intracellular staining a cytofix/cytoperm kit (BectonDickinson #554714) was used. After the labeling of the cell surface membrane proteins, cells were fixed for 10 minutes with the formaldehyde-based fixative solution, then washed and stained with primary and secondary (if required) antibodies diluted in the saponin-based washing buffer provided in the kit for 30 min to overnight. For detection of apoptotic cells the BD Annexin V kit (BD #560930) was used following the provided protocol. BrdU labeling was used to investigate *in vivo* proliferation of primary thymocytes, followed by detection of incorporated BrdU using the BD BrdU Flow Kit (#552598). Mice were injected i.p. with 150μl of a BrdU solution (10 mg/ml). After 2 hours, mice were sacrificed and thymi were collected. Membrane markers of thymocytes were labeled followed by fixation, permeabilization, DNAse treatment and anti-BrdU staining. Labeled cells were washed in PBS+1%BSA and acquired on a FACS Canto II (BD). Data was recorded in a low flow rate mode to ensure a good resolution of the labeling. Analyses were performed using FlowJo software (BD FlowJo LLC).

### TIRF Microscopy

Pooled thymocyte suspensions from 2-3 WT or L19A mice were obtained and LIN-thymocytes were negative selected using Ly6c-, F4/80-, CD11b-, CD19-, B220-, NK1.1-, CD4 - and CD8-specific antibodies and subsequently sheep-anti-rat dynabeads (Invitrogen #11035). A second round of selection using a FACSAria Fusion cell sorter was performed on LIN-thymocytes to obtain DN CD44-thymocytes (DN3+DN4 populations) that express GFP levels under 10^3^ relative fluorescence units. This latter selection criterion, excluding the ~1% brightest DN cells, guaranteed that we could detect individual particles at the cell surface using the TIRF setting. Sorted DN CD44-thymocytes were left to recover in RPMI containing 5% FBS for at least 2 hours. 30 minutes before TIRFM analysis, thymocytes were seeded on microscopy plates pretreated with fibronectin (10 μg/ml, 60 min, 37°C). Experiments were performed using a TIRF microscope (Leica AM TIRF inverted) equipped with an EM-CCD camera (Andor DU 885-CS0-#10-VP), a 100x oil-immersion objective (HCX PL APO 100x/1.46 NA) and a 488-nm diode laser. The microscope was equipped with incubator and temperature control units; experiments were performed at 37°C with 5% CO2. To minimize photobleaching effects before image acquisition, cells were located and focused using the bright field, and a fine focus adjustment in TIRF mode was made at 5% laser power, an intensity insufficient for single-particle detection that ensures negligible photobleaching. Image sequences of individual particles (500 frames) were acquired at 49% laser power with a frame rate of 10 Hz. The penetration depth of the evanescent field used was 90 nm. For the analysis, particles were detected and tracked using previously described algorithms (U-Track2; (Jaqaman et al., 2008)) implemented in MATLAB. The intensity value for each particle was calculated by subtracting in each frame the background intensity from the particle intensity. In addition, to minimize photon fluctuations within a given frame, the particle intensity was represented as the average value (background subtracted) obtained over the first 20 frames. For the short-time lag diffusion coefficient (D_1-4_), only tracks longer than 50 frames were used for further analysis; particles that merged or split were excluded. Individual trajectories were used to generate mean-square-displacement (MSD) plots that were required to extract the D_1-_ 4 using the equation: *MSD* = 4*D*1_4*t* + Δ0, where the Δ0 is the MSD offset at zero time lag. Additional details can be found in (Martinez-Munoz et al., 2018).

### Analysis of TCRβ repertoire

DN3a (CD4^-^CD8^-^CD25^+^CD44^-^) and blasting DP (CD4^+^CD8^+^CD69^-^FCS^hi^) thymocytes of WT and L19A 5-8 week-old male mice were purified by sorting. mRNA was extracted from purified thymocytes using the RNAeasy micro Kit (QIAGEN #74004). *Tcrb* amplicons were prepared using a 5’RACE-based protocol with the SMARTer Mouse TCR α/β Profiling Kit (Takara #634402) following the manufacturer’s instructions. The *Tcrb* amplicons were purified using AmpureXP beads (Beckman Coulter #A63880) and sequenced in parallel via Illumina High Throughput Sequencing on an Illumina MiSeq (2×300nt) at the Centro Nacional de Análisis Genómico (Barcelona, Spain). The quality of reads was assessed using FastQC (http://www.bioinformatics.babraham.ac.uk/projects/fastqc/). To extract CDR3 sequences, to determine V, D and J genes and to assemble clonotypes by CDR3 sequences, the raw sequencing reads were processed using the MiXCR v3.0.8 software (Bolotin et al., 2015). Further analysis of clones was conducted using the iNEXT R package (Chao et al., 2014), in-house written R-scripts and VDJtools v1.2.2 (Shugay et al., 2015). The ^*q*^*D* was computed using a rarefaction/extrapolation algorithm and coverage was estimated using the same techniques.

### Statistical analysis

Statistical parameters including the exact value of n, the means ± SD or SEM are described in the text and figure legends. All results were analyzed using GraphPad PRISM 7.0 (*p<0.05, ** p<0.01, *** p<0.001; **** p<0.0001 or the exact p value). TIRFM-Lag diffusion coefficient (D_1-4_) and the mean fluorescent intensity of single particles were analyzed using a two-tailed Mann-Whitney nonparametric test. All other statistical analyses were performed using the parametric, two-tailed and unpaired t-test or the Mann-Whitney test.

### Data availability

Raw sequencing data that underlie the analyses reported in figure 6 and supplementary figure 3 have been uploaded to the Bioproject Sequence Read Archive and can be obtained from the corresponding author upon reasonable request

## Materials Availability Statement

The mice generated in this study are available upon reasonable request from the Lead Contact with a completed Materials Transfer Agreement. Further information and requests for reagents may be directed to the corresponding author Dr. Hisse M. van Santen

## Acknowledgments

We thank Cristina Prieto, Valentina Blanco and Tania Gomez for excellent technical assistance and Belen Pintado and Veronica Dominguez from the CNB/CBMSO transgenic mouse facility for the generation of the CD3ζL19A-GFP and CD3ζWT-GFP mice. We thank Balbino Alarcon, Maria Luisa Toribio and our laboratory members for their expert advice during the development of this project and critical analysis of this manuscript. E.R.B. was supported by a FPI fellowship (BES-2014-068000, Subprograma Estatal de Formación del Programa Estatal de Promoción del Talento y su Empleabilidad, en el marco del Plan Estatal de Investigación Científica y Técnica y de Innovación 2013-2016) and an EMBO short term fellowship (#7272). This work was supported by Spanish Ministry of Economy and Competitiveness (MINECO) grants SAF2013-47975-R and SAF2016-76394-R to H.M.v.S. and FIS2016-78883-C2-2-P to M.C. The Centre for Genomic Regulation acknowledges support of the Spanish Ministry of Economy and Competitiveness, “Centro de Excelencia Severo Ochoa,” the Centres de Recerca de Catalunya Program/Generalitat de Catalunya. The Centro de Biología Molecular Severo Ochoa has been supported by the Fundación Ramón Areces.

## Author Contributions

E.R.B. and H.M.v.S. conceived the project. E.R.B. designed and conducted the experiments. E.M.G.C. conducted TIRF microscopy and performed analysis and interpretation of the obtained data. J.P., M.C. and D.A. analyzed and interpreted the TCRβ repertoire data, generating scripts necessary for visualization and interpretation of the results. M.M. provided ideas, designed experiments and helped with interpretation of results. E.R.B. and H.M.v.S. analyzed and interpreted the data and wrote the manuscript. All other authors edited the manuscript.

## Supplementary Material

Supplementary material consists of supplementary figures S1-S4

## Figure Legends

**Supplementary Figure 1. Expression of transgenic WT and L19A CD3ζ-GFP chains under control of a human CD2 promoter/enhancer cassette.** (a) Quantification of the total number of thymocytes in each mouse line. Each dot shows the total number of thymocytes in an individual mouse. Graphs presents the mean ± SEM. (b) Expression pattern of the transgenes. Thymocyte suspensions obtained from 6 week old WT (upper panels) and L19A (lower panels) mice were stained with fluorescently-labeled antibodies specific for the indicated proteins, acquired on a FACScanto flow cytometer and analyzed with FlowJo software. The WT and L19A CD3ζ-GFP chains are homogeneously expressed in all identified thymocyte populations. (c) Relative copy numbers of the transgenes. Genomic DNA isolated from tail snips from 5-6 mice of each of the transgenic lines was used as a template for a qPCR with oligonucleotides spanning the CD3ζ-GFP junction. For normalization, a qPCR amplifying the genomic sequence of the *Foxo1* gene was used. P-values were calculated using an unpaired two-tailed Student’s t test with 95% CI (***p<0.001).

**Supplementary Figure 2. Expression of the pre-TCR at the surface of DN thymocytes.** Expression levels of the transgenic (a) CD3ζ-GFP fusion proteins (detected via the GFP moiety) and CD3εγ and CD3εδ heterodimers (b) at the cell surface and (c) detected by intracellular staining (using the 2C11 mAb) for DN3a (Lin^-^CD44^-^ CD25^hi^), DN3b (Lin^-^CD44^-^CD25^lo^) and DN4 (Lin^-^CD44^-^CD25^-^) thymocytes from WT and L19A mice. Background levels of staining were determined using (a) *Rag1*-deficient (RAG^-/-^) or (b and c) *Cd3e*-deficient (Cd3e^-/-^) thymocytes. Graphs present the mean ± SEM of the expression level calculated for a total of 4 experiments. P-values were calculated using an unpaired two-tailed Student’s t test with 95% CI. (d) Correlation between levels of the GFP-coupled WT or L19A CD3ζ chains and cell surface labeling of CD3ε using the 2C11 antibody.

**Supplementary Figure 3. Properties of the TCRβ repertoire in DN and early DP thymocytes of WT and L19A mice.** (a) Number of clones per sample estimated with the ^*1*^*D* for WT (black) and L19A (red) DP thymocytes. Solid lines: Rarefaction (interpolation) curve; dashed lines: extrapolation curves; symbols: observed diversity. (b) Number of clones per sample estimated with the ^*2*^*D* for WT (black) and L19A (red) DP thymocytes. Solid lines: Rarefaction (interpolation) curve; dashed lines: extrapolation curves; symbols: observed diversity. (c) The coverage of the sampling process, that estimates the fraction of the real number of clones actually detected in the experiment for ^*0*^*D*. Solid lines: rarefaction (interpolated) coverage; Dashed lines: extrapolated coverage; Symbols: computed coverage using the actual experimental data. It can be seen that the coverage is almost 1 in all cases (the symbols are close to the asymptotic 100% coverage), meaning that the sampling has covered the real underlying repertoire almost completely. (d) Length distribution of the CDR3 region of WT and L19A DN3 and early DP thymocytes. Significant differences in distribution were not observed. (e) Scatter plot of frequencies of VDJ combinations observed in the repertoires of WT and L19A early DP thymocytes. The coordinates of each dot correspond to the mean of frequencies of each VDJ combination in WT and L19A early DP thymocytes. No statistically significant differences were observed for any of the combinations.

**Supplementary Figure 4. General gating strategy for DN thymocytes.** (a) Numbers in the plots indicate the gating order. LIN^-^ (CD4^-^, CD8^-^, CD19^-^; B220^-^ and CD11c^-^) thymocytes, panel 3, were gated using 4a or 4b strategy depending of the parameter to analyze.

## References

Blanco, R., Borroto, A., Schamel, W., Pereira, P., & Alarcon, B. (2014). Conformational changes in the T cell receptor differentially determine T cell subset development in mice. Sci Signal, 7(354), ra115. doi:10.1126/scisignal.2005650

Bolotin, D. A., Poslavsky, S., Mitrophanov, I., Shugay, M., Mamedov, I. Z., Putintseva, E. V., & Chudakov, D. M. (2015). MiXCR: software for comprehensive adaptive immunity profiling. Nat Methods, 12(5), 380–381. doi:10.1038/nmeth.3364

Borroto, A., Arellano, I., Dopfer, E. P., Prouza, M., Suchanek, M., Fuentes, M., … Alarcon, B. (2013). Nck recruitment to the TCR required for ZAP70 activation during thymic development. J Immunol, 190(3), 1103–1112. doi:10.4049/jimmunol.1202055

Borroto, A., Mallabiabarrena, A., Albar, J. P., Martinez, A. C., & Alarcon, B. (1998). Characterization of the region involved in CD3 pairwise interactions within the T cell receptor complex. J Biol Chem, 273(21), 12807–12816.

Brameshuber, M., Kellner, F., Rossboth, B. K., Ta, H., Alge, K., Sevcsik, E., … Huppa, J. B. (2018). Monomeric TCRs drive T cell antigen recognition. Nat Immunol, 19(5), 487–496. doi:10.1038/s41590-018-0092-4

Brodeur, J. F., Li, S., da Silva Martins, M., Larose, L., & Dave, V. P. (2009). Critical and multiple roles for the CD3epsilon intracytoplasmic tail in double negative to double positive thymocyte differentiation. J Immunol, 182(8), 4844–4853. doi:10.4049/jimmunol.0803679

Carrasco, Y. R., Navarro, M. N., & Toribio, M. L. (2003). A role for the cytoplasmic tail of the pre-T cell receptor (TCR) alpha chain in promoting constitutive internalization and degradation of the pre-TCR. J Biol Chem, 278(16), 14507–14513. doi:10.1074/jbc.M204944200

Chao, A., Gotelli, N. J., Hsieh, T. C., Sander, E. L., Ma, K. H., Colwell, R. K., & Ellison, A. M. (2014). Rarefaction and extrapolation with Hill numbers: a framework for sampling and estimation in species diversity studies. Ecological Monographs, 84(1), 45–67. doi:10.1890/13-0133.1

Cheng, A. M., Negishi, I., Anderson, S. J., Chan, A. C., Bolen, J., Loh, D. Y., & Pawson, T. (1997). The Syk and ZAP-70 SH2-containing tyrosine kinases are implicated in pre-T cell receptor signaling. Proc Natl Acad Sci U S A, 94(18), 9797–9801. doi:10.1073/pnas.94.18.9797

Das, D. K., Mallis, R. J., Duke-Cohan, J. S., Hussey, R. E., Tetteh, P. W., Hilton, M., … Reinherz, E. L. (2016). Pre-T Cell Receptors (Pre-TCRs) Leverage Vbeta Complementarity Determining Regions (CDRs) and Hydrophobic Patch in Mechanosensing Thymic Self-ligands. J Biol Chem, 291(49), 25292–25305. doi:10.1074/jbc.M116.752865

DeJarnette, J. B., Sommers, C. L., Huang, K., Woodside, K. J., Emmons, R., Katz, K., … Love, P. E. (1998). Specific requirement for CD3epsilon in T cell development. Proc Natl Acad Sci U S A, 95(25), 14909–14914.

Fahmy, T. M., Bieler, J. G., Edidin, M., & Schneck, J. P. (2001). Increased TCR avidity after T cell activation: a mechanism for sensing low-density antigen. Immunity, 14(2), 135–143.

Fantini, J., Di Scala, C., Baier, C. J., & Barrantes, F. J. (2016). Molecular mechanisms of protein-cholesterol interactions in plasma membranes: Functional distinction between topological (tilted) and consensus (CARC/CRAC) domains. Chem Phys Lipids, 199, 52–60. doi:10.1016/j.chemphyslip.2016.02.009

Gagnon, E., Schubert, D. A., Gordo, S., Chu, H. H., & Wucherpfennig, K. W. (2012). Local changes in lipid environment of TCR microclusters regulate membrane binding by the CD3epsilon cytoplasmic domain. J Exp Med, 209(13), 2423–2439. doi:10.1084/jem.20120790

Gascoigne, N. R., Rybakin, V., Acuto, O., & Brzostek, J. (2016). TCR Signal Strength and T Cell Development. Annu Rev Cell Dev Biol, 32, 327–348. doi:10.1146/annurev-cellbio-111315-125324

Gil, D., Schamel, W. W., Montoya, M., Sanchez-Madrid, F., & Alarcon, B. (2002). Recruitment of Nck by CD3 epsilon reveals a ligand-induced conformational change essential for T cell receptor signaling and synapse formation. Cell, 109(7), 901–912.

Godfrey, D. I., Kennedy, J., Suda, T., & Zlotnik, A. (1993). A developmental pathway involving four phenotypically and functionally distinct subsets of CD3-CD4-CD8-triple-negative adult mouse thymocytes defined by CD44 and CD25 expression. J Immunol, 150(10), 4244–4252.

Greaves, D. R., Wilson, F. D., Lang, G., & Kioussis, D. (1989). Human CD2 3’-flanking sequences confer high-level, T cell-specific, position-independent gene expression in transgenic mice. Cell, 56(6), 979–986.

Groettrup, M., Ungewiss, K., Azogui, O., Palacios, R., Owen, M. J., Hayday, A. C., & von Boehmer, H. (1993). A novel disulfide-linked heterodimer on pre-T cells consists of the T cell receptor beta chain and a 33 kd glycoprotein. Cell, 75(2), 283–294.

Hoffman, E. S., Passoni, L., Crompton, T., Leu, T. M., Schatz, D. G., Koff, A., … Hayday, A. C. (1996). Productive T-cell receptor beta-chain gene rearrangement: coincident regulation of cell cycle and clonality during development in vivo. Genes Dev, 10(8), 948–962. doi:10.1101/gad.10.8.948

Hogquist, K. A., & Jameson, S. C. (2014). The self-obsession of T cells: how TCR signaling thresholds affect fate ‘decisions’ and effector function. Nat Immunol, 15(9), 815–823. doi:10.1038/ni.2938

Irving, B. A., Alt, F. W., & Killeen, N. (1998). Thymocyte development in the absence of pre-T cell receptor extracellular immunoglobulin domains. Science, 280(5365), 905–908.

Ishikawa, E., Miyake, Y., Hara, H., Saito, T., & Yamasaki, S. (2010). Germ-line elimination of electric charge on pre-T-cell receptor (TCR) impairs autonomous signaling for beta-selection and TCR repertoire formation. Proc Natl Acad Sci U S A, 107(46), 19979–19984. doi:10.1073/pnas.1011228107

Jacobs, H., Vandeputte, D., Tolkamp, L., de Vries, E., Borst, J., & Berns, A. (1994). CD3 components at the surface of pro-T cells can mediate pre-T cell development in vivo. Eur J Immunol, 24(4), 934–939. doi:10.1002/eji.1830240423

Jaqaman, K., Loerke, D., Mettlen, M., Kuwata, H., Grinstein, S., Schmid, S. L., & Danuser, G. (2008). Robust single-particle tracking in live-cell time-lapse sequences. Nat Methods, 5(8), 695–702. doi:10.1038/nmeth.1237

Kumar, R., Ferez, M., Swamy, M., Arechaga, I., Rejas, M. T., Valpuesta, J. M., … van Santen, H. M. (2011). Increased sensitivity of antigen-experienced T cells through the enrichment of oligomeric T cell receptor complexes. Immunity, 35(3), 375–387.

Levelt, C. N., Mombaerts, P., Iglesias, A., Tonegawa, S., & Eichmann, K. (1993). Restoration of early thymocyte differentiation in T-cell receptor beta-chain-deficient mutant mice by transmembrane signaling through CD3 epsilon. Proc Natl Acad Sci U S A, 90(23), 11401–11405.

Lillemeier, B. F., Mortelmaier, M. A., Forstner, M. B., Huppa, J. B., Groves, J. T., & Davis, M. M. (2010). TCR and Lat are expressed on separate protein islands on T cell membranes and concatenate during activation. Nat Immunol, 11(1), 90–96.

Love, P. E., Shores, E. W., Johnson, M. D., Tremblay, M. L., Lee, E. J., Grinberg, A., … Westphal, H. (1993). T cell development in mice that lack the zeta chain of the T cell antigen receptor complex. Science, 261(5123), 918–921.

Mallis, R. J., Bai, K., Arthanari, H., Hussey, R. E., Handley, M., Li, Z., … Reinherz, E. L. (2015). Pre-TCR ligand binding impacts thymocyte development before alphabetaTCR expression. Proc Natl Acad Sci U S A, 112(27), 8373–8378. doi:10.1073/pnas.1504971112

Martin-Blanco, N., Blanco, R., Alda-Catalinas, C., Bovolenta, E. R., Oeste, C. L., Palmer, E., … Alarcon, B. (2018). A window of opportunity for cooperativity in the T Cell Receptor. Nat Commun, 9(1), 2618. doi:10.1038/s41467-018-05050-6

Martinez-Martin, N., Risueno, R. M., Morreale, A., Zaldivar, I., Fernandez-Arenas, E., Herranz, F., … Alarcon, B. (2009). Cooperativity between T cell receptor complexes revealed by conformational mutants of CD3epsilon. Sci Signal, 2(83), ra43.

Martinez-Munoz, L., Rodriguez-Frade, J. M., Barroso, R., Sorzano, C. O. S., Torreno-Pina, J. A., Santiago, C. A., … Mellado, M. (2018). Separating Actin-Dependent Chemokine Receptor Nanoclustering from Dimerization Indicates a Role for Clustering in CXCR4 Signaling and Function. Mol Cell, 71(5), 873. doi:10.1016/j.molcel.2018.08.012

Merkenschlager, M., Graf, D., Lovatt, M., Bommhardt, U., Zamoyska, R., & Fisher, A. G. (1997). How many thymocytes audition for selection? J Exp Med, 186(7), 1149–1158. doi:10.1084/jem.186.7.1149

Mingueneau, M., Kreslavsky, T., Gray, D., Heng, T., Cruse, R., Ericson, J., … Turley, S. (2013). The transcriptional landscape of alphabeta T cell differentiation. Nat Immunol, 14(6), 619–632. doi:10.1038/ni.2590

Minguet, S., Swamy, M., Alarcon, B., Luescher, I. F., & Schamel, W. W. (2007). Full activation of the T cell receptor requires both clustering and conformational changes at CD3. Immunity., 26(1), 43-54. Epub 2006 Dec 2021.

Molina, T. J., Kishihara, K., Siderovski, D. P., van Ewijk, W., Narendran, A., Timms, E., … et al. (1992). Profound block in thymocyte development in mice lacking p56lck. Nature, 357(6374), 161–164. doi:10.1038/357161a0

Molnar, E., Swamy, M., Holzer, M., Beck-Garcia, K., Worch, R., Thiele, C., … Schamel, W. W. (2012). Cholesterol and sphingomyelin drive ligand-independent T-cell antigen receptor nanoclustering. J Biol Chem, 287(51), 42664–42674. doi:10.1074/jbc.M112.386045

Mombaerts, P., Iacomini, J., Johnson, R. S., Herrup, K., Tonegawa, S., & Papaioannou, V. E. (1992). RAG-1-deficient mice have no mature B and T lymphocytes. Cell, 68(5), 869–877. doi:10.1016/0092-8674(92)90030-g

Pang, S. S., Berry, R., Chen, Z., Kjer-Nielsen, L., Perugini, M. A., King, G. F., … Rossjohn, J. (2010). The structural basis for autonomous dimerization of the pre-T-cell antigen receptor. Nature, 467(7317), 844–848. doi:10.1038/nature09448

Panigada, M., Porcellini, S., Barbier, E., Hoeflinger, S., Cazenave, P. A., Gu, H., … Grassi, F. (2002). Constitutive endocytosis and degradation of the pre-T cell receptor. J Exp Med, 195(12), 1585–1597. doi:10.1084/jem.20020047

Pivniouk, V., Tsitsikov, E., Swinton, P., Rathbun, G., Alt, F. W., & Geha, R. S. (1998). Impaired viability and profound block in thymocyte development in mice lacking the adaptor protein SLP-76. Cell, 94(2), 229–238.

Risueno, R. M., Gil, D., Fernandez, E., Sanchez-Madrid, F., & Alarcon, B. (2005). Ligand-induced conformational change in the T-cell receptor associated with productive immune synapses. Blood, 106(2), 601-608. Epub 2005 Mar 2024.

Rossboth, B., Arnold, A. M., Ta, H., Platzer, R., Kellner, F., Huppa, J. B., … Schutz, G. J. (2018). TCRs are randomly distributed on the plasma membrane of resting antigen-experienced T cells. Nat Immunol, 19(8), 821–827. doi:10.1038/s41590-018-0162-7

Saint-Ruf, C., Panigada, M., Azogui, O., Debey, P., von Boehmer, H., & Grassi, F. (2000). Different initiation of pre-TCR and gammadeltaTCR signalling. Nature, 406(6795), 524–527. doi:10.1038/35020093

Schamel, W. W., Arechaga, I., Risueno, R. M., van Santen, H. M., Cabezas, P., Risco, C., … Alarcon, B. (2005). Coexistence of multivalent and monovalent TCRs explains high sensitivity and wide range of response. J Exp Med, 202(4), 493-503. Epub 2005 Aug 2008.

Shah, D. K., & Zuniga-Pflucker, J. C. (2014). An overview of the intrathymic intricacies of T cell development. J Immunol, 192(9), 4017–4023. doi:10.4049/jimmunol.1302259

Shendure, J., & Ji, H. (2008). Next-generation DNA sequencing. Nat Biotechnol, 26(10), 1135–1145. doi:10.1038/nbt1486

Sherman, E., Barr, V., Manley, S., Patterson, G., Balagopalan, L., Akpan, I., … Samelson, L. E. (2011). Functional nanoscale organization of signaling molecules downstream of the T cell antigen receptor. Immunity, 35(5), 705–720.

Shugay, M., Bagaev, D. V., Turchaninova, M. A., Bolotin, D. A., Britanova, O. V., Putintseva, E. V., … Chudakov, D. M. (2015). VDJtools: Unifying Post-analysis of T Cell Receptor Repertoires. PLoS Comput Biol, 11(11), e1004503. doi:10.1371/journal.pcbi.1004503

Taghon, T., Yui, M. A., Pant, R., Diamond, R. A., & Rothenberg, E. V. (2006). Developmental and molecular characterization of emerging beta- and gammadelta-selected pre-T cells in the adult mouse thymus. Immunity, 24(1), 53–64. doi:10.1016/j.immuni.2005.11.012

van Meerwijk, J. P., Marguerat, S., Lees, R. K., Germain, R. N., Fowlkes, B. J., & MacDonald, H. R. (1997). Quantitative impact of thymic clonal deletion on the T cell repertoire. J Exp Med, 185(3), 377–383. doi:10.1084/jem.185.3.377

Viret, C., & Janeway, C. A., Jr. (1999). MHC and T cell development. Rev Immunogenet, 1(1), 91–104.

von Boehmer, H. (2005). Unique features of the pre-T-cell receptor alpha-chain: not just a surrogate. Nat Rev Immunol, 5(7), 571–577. doi:10.1038/nri1636

Wang, F., Beck-Garcia, K., Zorzin, C., Schamel, W. W., & Davis, M. M. (2016). Inhibition of T cell receptor signaling by cholesterol sulfate, a naturally occurring derivative of membrane cholesterol. Nat Immunol, 17(7), 844–850. doi:10.1038/ni.3462

Wang, Y., Becker, D., Vass, T., White, J., Marrack, P., & Kappler, J. W. (2009). A conserved CXXC motif in CD3epsilon is critical for T cell development and TCR signaling. PLoS Biol, 7(12), e1000253. doi:10.1371/journal.pbio.1000253

Williams, J. A., Hathcock, K. S., Klug, D., Harada, Y., Choudhury, B., Allison, J. P., … Hodes, R. J. (2005). Regulated costimulation in the thymus is critical for T cell development: dysregulated CD28 costimulation can bypass the pre-TCR checkpoint. J Immunol, 175(7), 4199–4207. doi:10.4049/jimmunol.175.7.4199

Xu, C., Gagnon, E., Call, M. E., Schnell, J. R., Schwieters, C. D., Carman, C. V., … Wucherpfennig, K. W. (2008). Regulation of T cell receptor activation by dynamic membrane binding of the CD3epsilon cytoplasmic tyrosine-based motif. Cell, 135(4), 702–713. doi:10.1016/j.cell.2008.09.044

Yamasaki, S., Ishikawa, E., Sakuma, M., Ogata, K., Sakata-Sogawa, K., Hiroshima, M., … Saito, T. (2006). Mechanistic basis of pre-T cell receptor-mediated autonomous signaling critical for thymocyte development. Nat Immunol, 7(1), 67–75.

Zhang, W., Sommers, C. L., Burshtyn, D. N., Stebbins, C. C., DeJarnette, J. B., Trible, R. P., … Samelson, L. E. (1999). Essential role of LAT in T cell development. Immunity, 10(3), 323–332.

Zhong, L., Zeng, G., Lu, X., Wang, R. C., Gong, G., Yan, L., … Chen, Z. W. (2009). NSOM/QD-based direct visualization of CD3-induced and CD28-enhanced nanospatial coclustering of TCR and coreceptor in nanodomains in T cell activation. PLoS One, 4(6), e5945. doi:10.1371/journal.pone.0005945

