## Supplementary material for "Pre-TCR signaling intensity shapes the TCRβ repertoire": suppl_figs_1_4

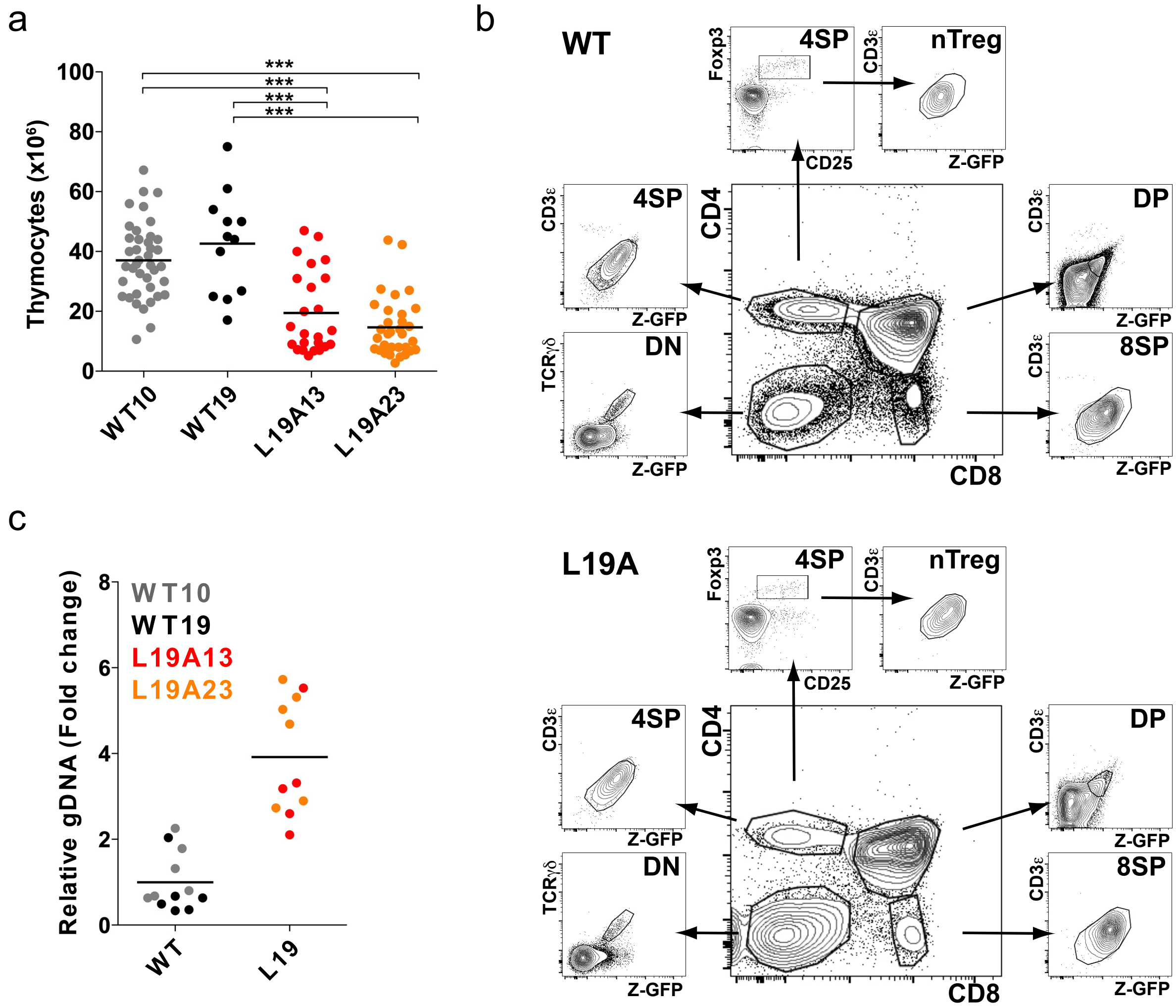


*Bovolenta et al., Figure Sup 1*

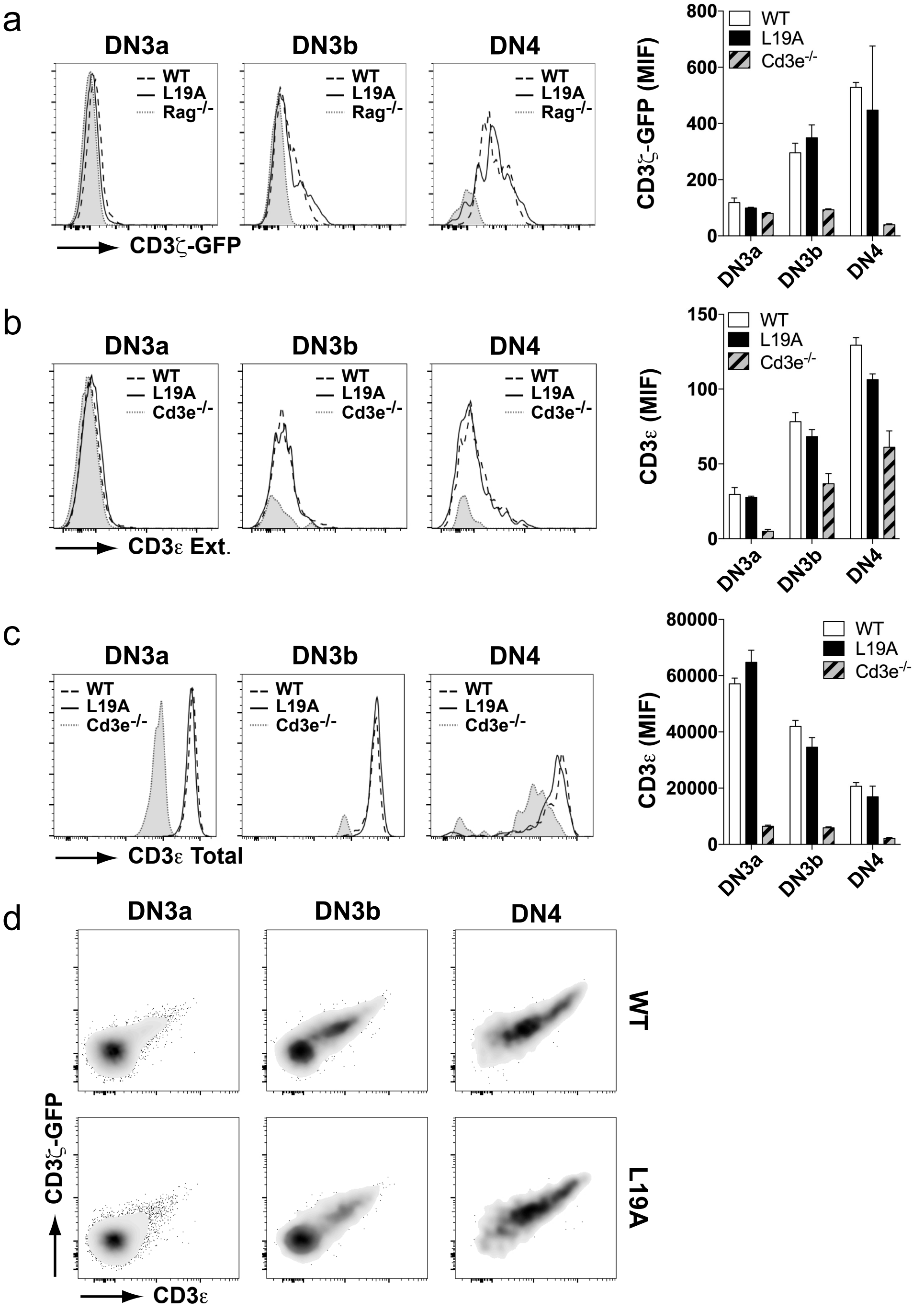


*Bovolenta et al., Figure Sup 2*

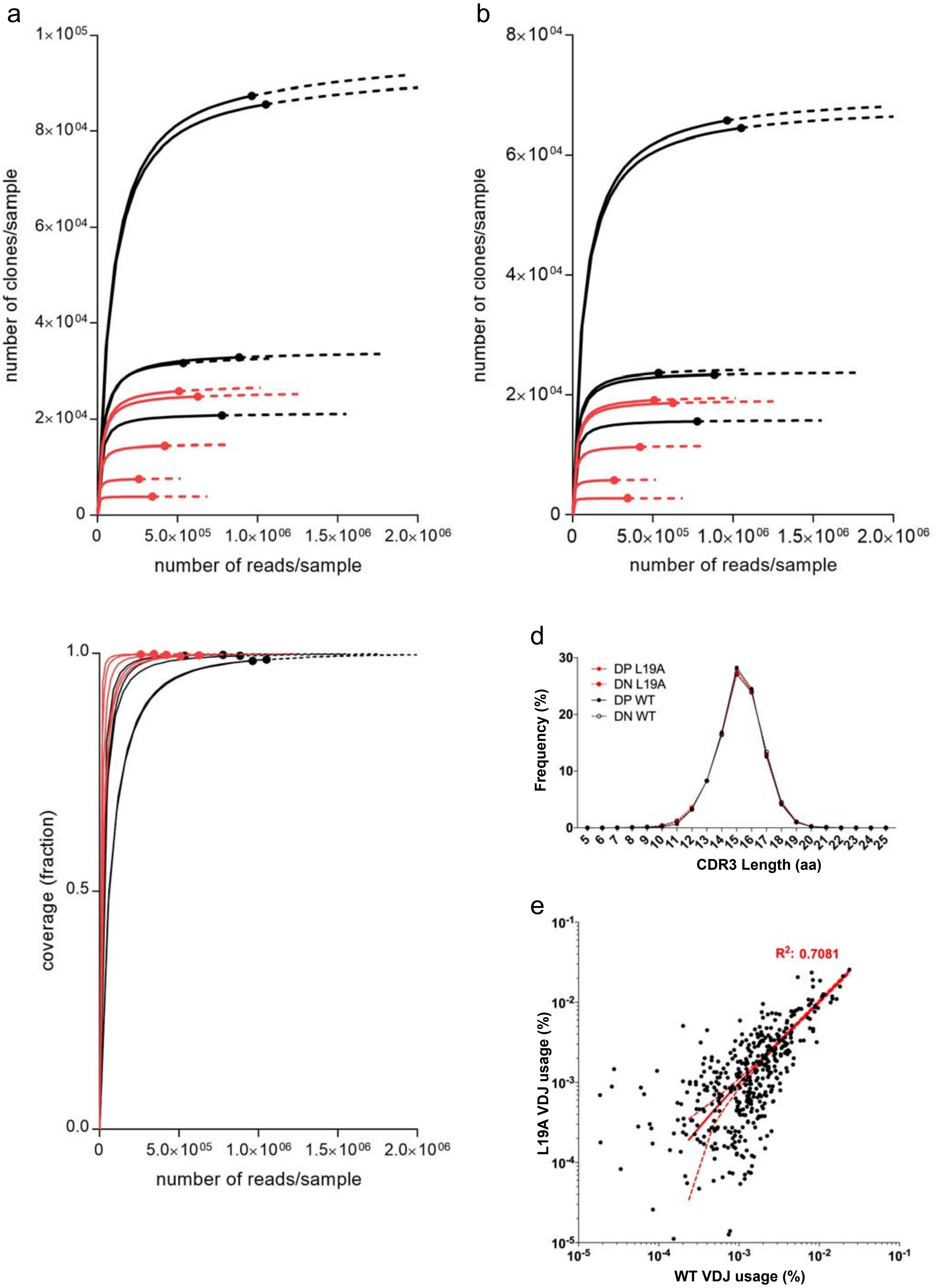


*Bovolenta et al., Suppl Figure 3*

a

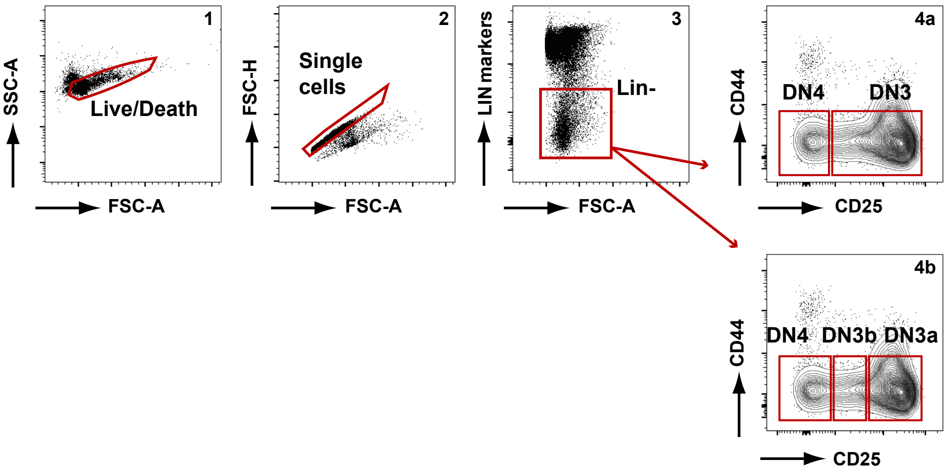


*Bovolenta et al., Figure Sup 4*
